# Bayesian Factor Analysis for Binary and Ordinal Phenotypes with Missingness

**DOI:** 10.64898/2026.07.15.733660

**Authors:** N. Shashaank, David A. Knowles

## Abstract

Binary and ordinal phenotypes are common in clinical screening and self-reported questionnaires, but many factor analysis and matrix factorization methods are only applicable for quantitative phenotypes with real-valued and/or continuous data distributions. To address this, we propose FABOr (Factor Analysis of Binary and Ordinal data), a Bayesian framework for matrix factorization in which the low-rank matrices are latent variables with continuous priors while the phenotypes are observed variables modeled with appropriate binary/ordinal likelihoods. We also develop missing not at random (MNAR) extensions of FABOr for analyzing data with structured missingness. In experiments with simulated phenotypes, we found that FABOr performs similarly to the best-performing benchmark methods on binary data at imputation and exceeds the performance of all tested benchmark methods on ordinal data. We then applied FABOr to analyze real-world binary and ordinal phenotypes from the Simons Foundation SPARK dataset on autism spectrum disorder (ASD) and found that it improved imputation accuracy by up to 5% on binary data and up to 23% on ordinal data relative to the benchmark methods.

## 1 Introduction

Clinical questionnaires and surveys often contain phenotypes that are binary (e.g., 0 or 1, yes or no) or ordinal, which take on an ordered set of values (e.g., low, medium, high). Binary and ordinal variables are used to represent important medical phenotypes such as disease status/severity rating and self-reported phenotypes such as questions about an individual’s physical/mental health and well-being. However, classical factor analysis [1] and matrix factorization [2, 3] algorithms for identifying the relationships and latent factors underlying multiple variables are only applicable to quantitative phenotypes (i.e., real-valued and/or continuous data). To address this, some works adapt existing factor analysis methods and apply a transformation to the continuous-valued latent factors to compare against binary or ordinal data [4–6], while others design new methods that constrain the latent factors to be discrete-valued as well [7–9]. Many factor analysis methods are also not equipped to handle missing not at random (MNAR) data, where certain questions are left blank by the individual respondents depending on the questionnaire structure or for undisclosed reasons that may depend on their unobserved state.

Here, we present Factor Analysis of Binary and Ordinal data (FABOr), a Bayesian framework for matrix factorization where the low-rank matrices are treated as latent variables (Fig. 1). In contrast to classical matrix factorization where the product of the low-rank matrices is optimized to directly approximate the data, FABOr approximates the posterior distribution over the matrices given appropriate priors and the data likelihood. This allows the binary/ordinal phenotypes to be modeled with the appropriate constraints while the underlying factors are modeled more flexibly within the real-valued or continuous latent space depending on the chosen priors. FABOr also includes extensions for modeling MNAR data by using the latent variables to model both the phenotypes’ observed values and the missingness pattern.

**Fig. 1:**
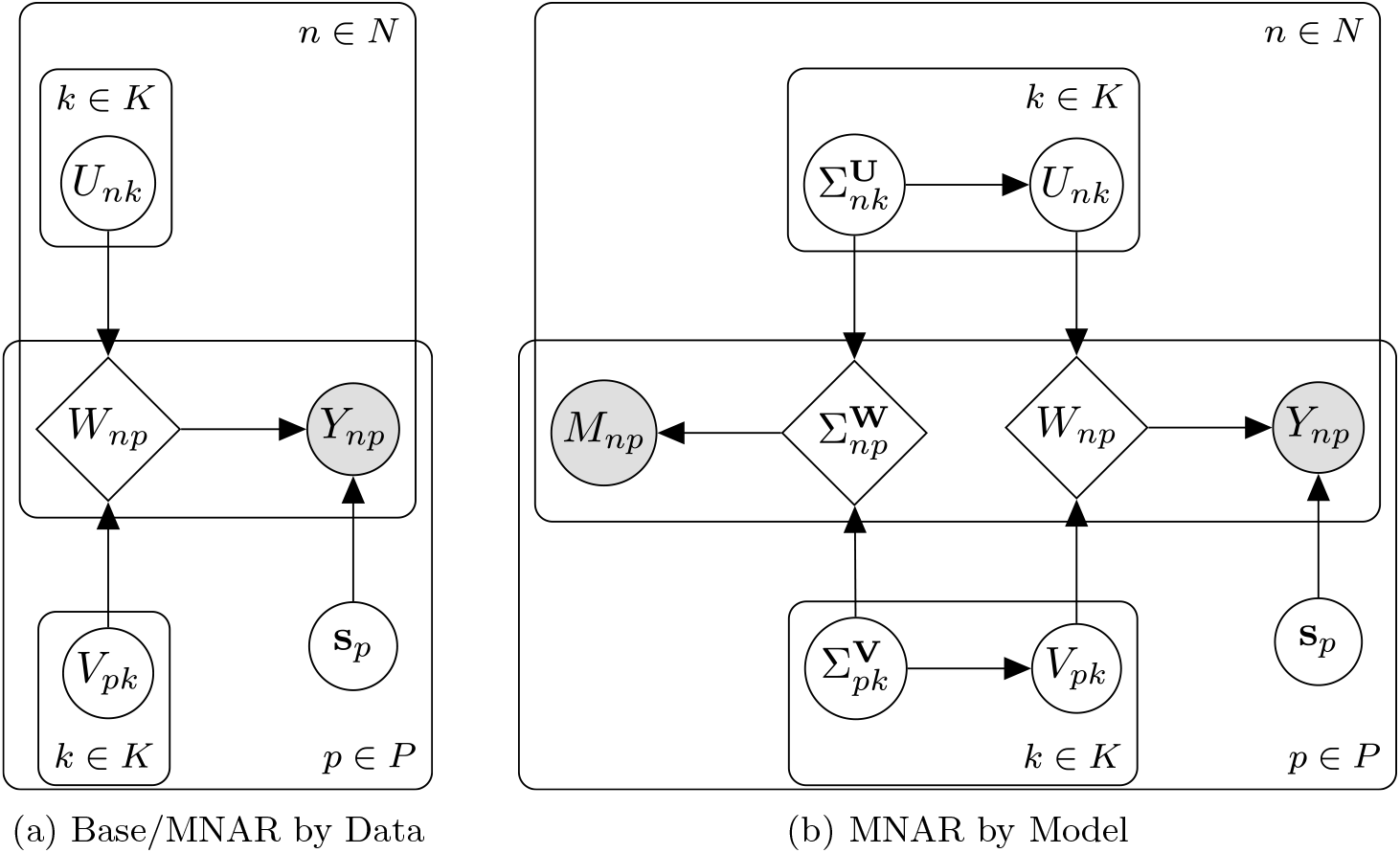
Probabilistic graphical models underlying FABOr. In the base model and MNAR by Data extension **(a)**, the low-rank matrices **U** and **V** are modeled as latent variables influencing the likelihood of the observed data **Y**, while in the MNAR by Model extension **(b)**, additional low-rank matrices **Σ**^**U**^ and **Σ**^**V**^ influence **U** and **V** based on the structure of the observed missingness pattern **M**. Latent variables are denoted as unshaded circles, deterministic variables (i.e., variables which are deterministic functions of other variables) are denoted as unshaded diamonds, and observed variables are denoted as shaded circles.

## 2 Methods

### 2.1 Factor Analysis Model

Let **Y** denote the dataset of interest, where each entry

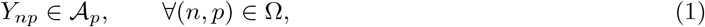

corresponds to one of the allowable observed values *A*_*p*_ for phenotype *p*, and Ω ⊆ {1, …, *N*}×{1, …, *P*} is the set of observed (non-missing) entries. Let *D*_*p*_ = |*A*_*p*_| be the number of categories for phenotype *p*, such that *D*_*p*_ = 2 for binary phenotypes and *D*_*p*_ > 2 for ordinal phenotypes. Our objective is to identify two lower-dimensional matrices **U** of size *N* × *K* and **V** of size *P* × *K* to approximate the distribution of responses in **Y** using a small number of latent factors *K* ≪*P* .

Under the base probabilistic framework in FABOr (Fig. 1a), **Y** is treated as an observed variable whose likelihood function is parameterized by latent variables **U** and **V**. The likelihood function is determined by an intermediate matrix **W** of size *N* × *P* representing the transformed product of **U** and **V**.

In the simplest instantiation of this framework (which we call the *Normal model* ), **U** and **V** are real-valued matrices with zero-centered element-wise Gaussian priors,

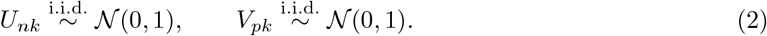

**W** is transformed into a valid probability matrix within the unit interval using a link function *f*,

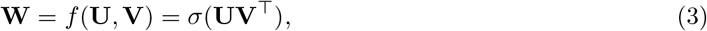

which for the real coordinate space can be achieved using the sigmoid function *σ*(*x*) = 1/(1 + exp(−*x*)) applied element-wise.

Binary phenotypes are modeled using a Bernoulli likelihood,

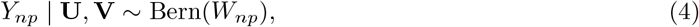

analogous to logistic regression. For ordinal phenotypes, we adapted an approach for ordinal regression [10]. Specifically, we probabilistically discretize *W*_*np*_ into the observed categories *A*_*p*_ using a series of cutpoints **c**_*p*_, which has the effect of inducing a natural ordering of the categories. We define **c**_*p*_ as a deterministic function of an additional per-phenotype latent variable 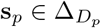,

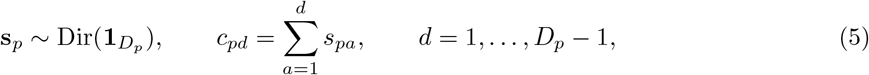

where Dir is a Dirichlet distribution. **c**_*p*_ and *W*_*np*_ are subsequently mapped from the unit interval to the real line using the inverse sigmoid (logit) function *σ*^−1^(*x*) = log(*x*/(1 − *x*)). The likelihood is a categorical distribution,

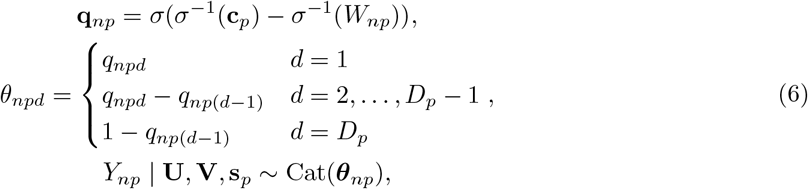

where the probability mass for each category *θ*_*npd*_ is determined from the cumulative probabilities **q**_*np*_, which are computed by subtracting the logit-transformed *W*_*np*_ from the logit-transformed **c**_*p*_. While we theoretically could have kept them both in the unit interval and applied a linear transformation when computing **q**_*np*_, we found that the non-linear transformation afforded by the sigmoid and logit functions performed better in practice, possibly due to a more favorable optimization landscape for variational inference.

We fit the FABOr model using stochastic variational inference (SVI) [11–13], a gradient-based inference method that approximates the posterior distribution using a variational distribution by maximizing the evidence lower bound (ELBO), composed of the joint log-likelihood and the entropy of the latent variables. The ELBO is only computed on observed entries Ω in the original data to prevent missing responses from affecting the posterior. For non-Dirichlet priors, we selected the mean-field Gaussian family for the variational distribution, while for Dirichlet priors we optimized within the simplex as point estimates using maximum a posteriori (MAP) inference.

Because the binary and ordinal likelihoods are modeled separately for each phenotype from the same probability matrix, FABOr supports mixing binary and ordinal phenotypes in the same dataset (including mixed ordinal scales), which is currently not supported by many factor analysis models.

### 2.2 Non-Gaussian Latent Factor Model Variants

**U** and **V** can be given any continuous-valued prior, as long as *f* can be constructed as a differentiable function that transforms the product into the unit interval (i.e., *f* : supp(**UV**^⊤^) → [0, 1]). To test whether non-Gaussian priors can improve the performance of FABOr, we created three different model variants.

In the *Dirichlet-Beta model*, **U** is drawn from a Dirichlet prior and **V** is drawn from a Beta prior with fixed hyperparameters *α* = 2 and *β* = 2,

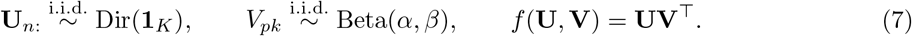

The Dirichlet prior on **U** enforces a *K*-dimensional simplex constraint such that *U*_*nk*_ ≥0 and ∑_*k*_ *U*_*nk*_ = 1, a probability vector over the latent factors. Subsequently, the loadings for each individual (i.e., the probability vector) can be used to interpret how they are affected by each latent factor based on their observed phenotypes. Similarly, the Beta prior on **V** enforces a unit interval constraint such that **V** ∈ [0, 1], which can be more meaningful when interpreting the loadings for each phenotype to understand which latent factors are more/less strongly associated with it.

In the *Dirichlet-Normal model*, we combine the Dirichet prior for **U** from the Dirichlet-Beta model with the Gaussian prior for **V** from the Normal model such that,

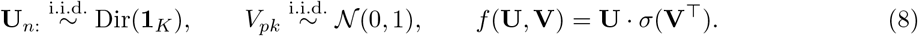

so that each **U**_*n*:_ belongs to the probability simplex but leaves **V** unbounded. It also serves to evaluate which prior for **U** (Gaussian or Dirichlet) and **V** (Gaussian or Beta) has a greater impact on the results by comparing against the other models.

In the *Lognormal model*, **U** and **V** are drawn from zero-centered Lognormal priors,

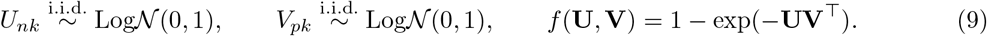

The Lognormal priors enforce non-negativity on the latent factors (i.e., *U*_*nk*_, *V*_*pk*_ ∈ [0, ∞ )), making the model analogous to non-negative matrix factorization. It can be used in place of the Normal model when the magnitudes of the latent factor loadings are more useful to interpret than their directions.

In all of these model variants, the likelihood of the binary/ordinal phenotypes remains the same because we only change how **W** is computed from **U** and **V**, thereby creating a consistent structure for comparison during training and evaluation.

### 2.3 Missing Not at Random Extensions

Baseline FABOr implicitly assumes that the data is missing at random (MAR). However, accounting for structured missingness that may be present in MNAR data requires modeling the missingness matrix **M** where *M*_*np*_ = 1 if (*n, p*) ∈ Ω and *M*_*np*_ = 0 otherwise; i.e., 1 denotes an observed element and 0 denotes a missing element. We tested two approaches for incorporating **M**. In the first approach, which we call “MNAR by Data”, **M** is modeled as an extension of the original data,

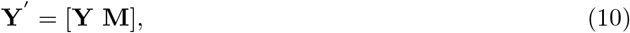

and the models are subsequently fit on **Y**′ instead of **Y**.

In the second approach, which we call “MNAR by Model”, **M** is modeled as a separate observed variable with a Bernoulli likelihood determined by two additional latent variables **Σ**^**U**^ and **Σ**^**V**^ (Fig. 1b), such that,

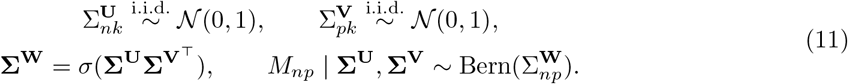

**Σ**^**U**^ and **Σ**^**V**^ are subsequently used to model the variance of **U** and **V**,

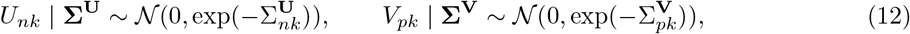

with lower variance corresponding to a lower probability that the element is missing and higher variance corresponding to a higher probability that the element is missing.

To avoid optimization difficulties due to the hierarchical relationship between **Σ**^**U**^ to **U** and **Σ**^**V**^ to **V**, we used non-centered parameterization [14] to split **U** and **V** into multiple independent variables,

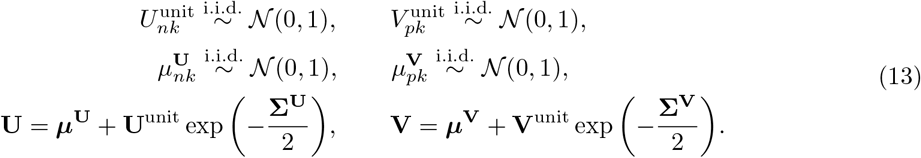

This construction is equivalent to Equation (12) but is easier to fit with the variational posterior on **U**^unit^ and **V**^unit^ rather than directly on **U** and **V**.

## 3 Experimental Setup

### 3.1 Datasets

We ran our experiments on simulated phenotypes generated with a latent factor model and real-world phenotypes taken from the Simons Foundation SPARK dataset on autism spectrum disorder (ASD).

#### 3.1.1 Simulated phenotypes

We generated two synthetic datasets: one with binary phenotypes and one with ordinal phenotypes. The base dimensions for both datasets were *N* = 100000, *P* = 100, and *K* = 6. The low-rank matrices were generated as,

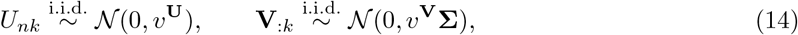

where each element of **U** is drawn from a zero-mean Gaussian distribution with variance *v*^**U**^ and each column of **V** is drawn from a zero-mean multivariate Gaussian distribution correlated across phenotypes with covariance *v*^**V**^. As questionnaires tend to have related questions ordered close to each other, the covariance matrix **Σ** follows an autoregressive structure,

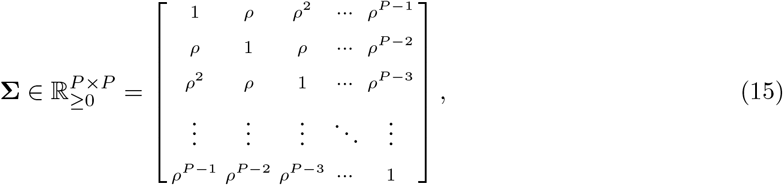

where *ρ* is a correlation coefficient. The values for *v*^**U**^, *v*^**V**^, and *ρ* were selected based on estimates of correlation in the real-world questionnaires we used from the SPARK dataset.

For the binary phenotype dataset, we set 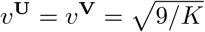 and *ρ* = 0.35. As phenotypes generally do not have an even distribution of 0s and 1s in real-world datasets, we used a per-column intercept **z** ∈ ℝ^*P*^ to offset the product of **U** and **V** when computing **W** with the sigmoid function to skew the distribution of values more towards 0 or 1 for each phenotype,

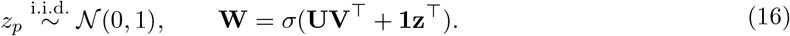

**Y** was drawn according to the binary likelihood in Equation (4).

For the ordinal phenotype dataset, we set 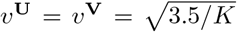 and *ρ* = 0.97. **W** was computed from the sigmoid function with no additional intercept according to Equation (3), and **Y** was drawn according to the ordinal likelihood in Equations (5)–(6). We set the number of categories *D*_*p*_ = 4 for all phenotypes.

To analyze the impact of missing entries on model performance, we created three different missingness patterns for each dataset. In the “Full” configuration, all elements are observed, i.e., *M*_*np*_ = 1. In the “MAR” configuration, each element has an 80% chance of being observed independent of phenotype or value as determined by a Bernoulli distribution, i.e., 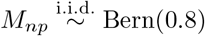. In the “MNAR” configuration, the probability of an element being observed is dependent on both the observed value and the phenotype,

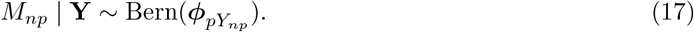

For some phenotypes, lower values are more likely to be observed, while for other phenotypes higher values are more likely to be observed; this direction is determined by a Bern(0.5) draw for each phenotype. If lower values are more likely to be observed, ***ϕ***_*p*_ = [0.95, 0.65] for binary phenotypes and ***ϕ***_*p*_ = [0.95, 0.875, 0.725, 0.65] for ordinal phenotypes. If higher values are more likely to be observed, ***ϕ***_*p*_ = [0.65, 0.95] for binary phenotypes and ***ϕ***_*p*_ = [0.65, 0.725, 0.875, 0.95] for ordinal phenotypes.

#### 3.1.2 Real-world phenotypes

The SPARK dataset [15] contains genotypes and phenotypes for ASD individuals and unaffected family members and is currently the largest dataset dedicated to ASD. For binary phenotypes, we used the Social Communication Questionnaire (SCQ), which measures verbal and non-verbal communication abilities. It contains *P* = 40 yes-or-no questions and was completed by *N* = 81088 individuals (both ASD cases and non-ASD controls), with ∼ 97% of entries observed. For ordinal phenotypes, we used the Repetitive Behavior Scale-Revised (RBS-R), which measures the frequency and severity of restrictive and/or repetitive behaviors. It contains *P* = 43 4-point Likert scale questions and was completed by *N* = 42308 individuals (only ASD cases), with ∼99.9% of entries observed.

### 3.2 Benchmark Models

For binary phenotypes, we compared FABOr against two models: matrix factorization with a logistic likelihood which we call Logistic MF, and *k*-Greedy binary matrix factorization (BMF) [9]. In Logistic MF, the latent factors are real-valued, similar to the Normal model from FABOr but without Bayesian priors, and optimized using stochastic gradient descent against a binary cross-entropy loss. In *k*-Greedy BMF, the latent factors are constrained to be binary and optimized using a greedy search algorithm.

For ordinal phenotypes, we compared FABOr against two models: matrix factorization with an ordinal likelihood which we call Ordinal MF, and ordinal non-negative matrix factorization (OrdNMF) [6]. In Ordinal MF, the latent factors are (once again) unconstrained real-valued without Bayesian priors and optimized using stochastic gradient descent against a categorical cross-entropy loss. In OrdNMF, the latent factors are constrained to be non-negative and optimized using SVI as part of a probabilistic model. Both models include learnable cutpoints defining the thresholds between ordinal categories in the continuous-valued coordinate space.

### 3.3 Model Training and Evaluation

For each dataset, we randomly split the observed entries Ω into train/test sets, using a distribution of 90% train/10% test. The train set was used to fit the models with the appropriate optimization method, after which we imputed the test set entries using the models’ predictions to evaluate the performance against the ground truth original data. To ensure that the missingness pattern generated by the test set does not affect the interpretation of the missingness pattern inherent to the dataset during training, we performed the 90%/10% split on each phenotype (i.e., each column) so that the missingness imposed by the test set is uniformly distributed. We ran 10 random restarts of each model, with up to 1000 iterations per restart, and averaged the performance across restarts. We also tested several values of *K* ∈ {3, …, 10} for each model to identify the optimal number of factors for fitting the model while retaining useful information. Our main evaluation metric on the held-out test set was accuracy (i.e., the percentage of predictions that were equal to the ground truth). For simulated phenotypes, we used the mean squared error (MSE) on **W** as a secondary metric to determine our models’ performance in recovering the true latent factors. We measured the runtime for each of our models on an NVIDIA RTX Pro 6000 Blackwell GPU.

### 3.4 Code

FABOr was implemented in Pyro [16] and is available as a Python package at https://github.com/daklab/fabor. For the benchmark models, Logistic MF and Ordinal MF were implemented in PyTorch [17], and [*k*-Greedy BMF](https://github.com/kovacsrekaagnes/rank_k_Binary_Matrix_Factorisation) and [Ordinal NMF](https://github.com/Oligou/OrdNMF) were converted from the original repos in NumPy to PyTorch. The code to reproduce the experiments is available in a separate repository at https://github.com/skresearcher/fabor-experiments.

## 4 Results

Table 1 shows the performance comparison across all models tested in our experiments for the best-performing value of *K* with simulated and real-world phenotypes. We used imputation accuracy as our primary evaluation metric to best determine whether the models are able to capture the true structure of the original data.

**Table 1:** Evaluation of factor analysis models on simulated and real-world phenotypes. The average accuracy across 10 random restarts is reported for the best performing value of *K*.

|  |  | Simulated Binary |  |  | Simulated Ordinal |  |  | Real-World |  |
| --- | --- | --- | --- | --- | --- | --- | --- | --- | --- |
|  |  | Full | MAR | MNAR | Full | MAR | MNAR | SCQ | RBS-R |
| Benchmarks | Logistic MF | 0.809 | 0.800 | 0.807 | – | – | – | 0.773 | – |
| | $k$-Greedy BMF [9] | 0.649 | 0.643 | 0.658 | – | – | – | 0.751 | – |
|  | Ordinal MF | – | – | – | 0.531 | 0.516 | 0.530 | – | 0.580 |
|  | OrdNMF [6] | – | – | – | 0.328 | 0.196 | 0.208 | – | 0.496 |
| FABOr<br>Base | Normal | 0.808 | 0.803 | 0.809 | **0.572** | **0.568** | 0.573 | 0.793 | **0.601** |
|  | Dirichlet-Beta | 0.758 | 0.753 | 0.762 | 0.558 | 0.553 | 0.559 | 0.780 | 0.582 |
|  | Dirichlet-Normal | 0.759 | 0.753 | 0.762 | 0.558 | 0.554 | 0.560 | 0.780 | 0.583 |
|  | Lognormal | 0.741 | 0.731 | 0.741 | 0.560 | 0.554 | 0.561 | 0.778 | 0.592 |
| FABOr<br>MNAR by Data | Normal | **0.811** | **0.805** | **0.812** | **0.572** | **0.568** | **0.574** | **0.798** | **0.601** |
|  | Dirichlet-Beta | 0.758 | 0.753 | 0.765 | 0.557 | 0.553 | 0.559 | 0.782 | 0.582 |
|  | Dirichlet-Normal | 0.758 | 0.753 | 0.765 | 0.558 | 0.553 | 0.560 | 0.782 | 0.582 |
|  | Lognormal | 0.743 | 0.733 | 0.747 | 0.561 | 0.556 | 0.563 | 0.779 | 0.591 |
| FABOr<br>MNAR by Model | Normal | 0.804 | 0.797 | 0.806 | 0.563 | 0.557 | 0.563 | 0.785 | 0.590 |

### 4.1 Simulated Phenotype Study

When comparing the base Normal model from FABOr with the benchmark models for simulated binary and ordinal phenotypes, FABOr performs ∼ 25% better than k-Greedy BMF, comparable to Logistic MF, 2 – 3x better than OrdNMF, and 7 – 9% better than Ordinal MF. This demonstrates that the simplest configuration of FABOr achieves similar to or better performance than the recent literature while accommodating both binary and ordinal phenotypes in the same model if needed (Fig. 2). Among the benchmark models, the classical matrix factorization models with binary/ordinal likelihoods (Logistic MF and Ordinal MF) perform much better than the specialized methods for binary or ordinal matrix factorization (*k*-Greedy BMF and OrdNMF), which suggests that basic matrix factorization is a good starting point for binary/ordinal factor analysis and validates our approach with FABOr. When comparing the performance of these models across different values of *K*, the same observations hold: the Normal model from FABOr performs comparably to Logistic MF and outperforms *k*-Greedy BMF, Ordinal MF, and OrdNMF on all values of *K* tested (Fig. S1). On the binary dataset, the Normal model achieves the highest accuracy at *K* = 7, which is close to the true number of latent factors in the dataset (*K* = 6). Similarly, on the ordinal dataset the Normal model achieves the highest accuracy at *K* = 6, which matches the true number of latent factors.

**Fig. 2:**
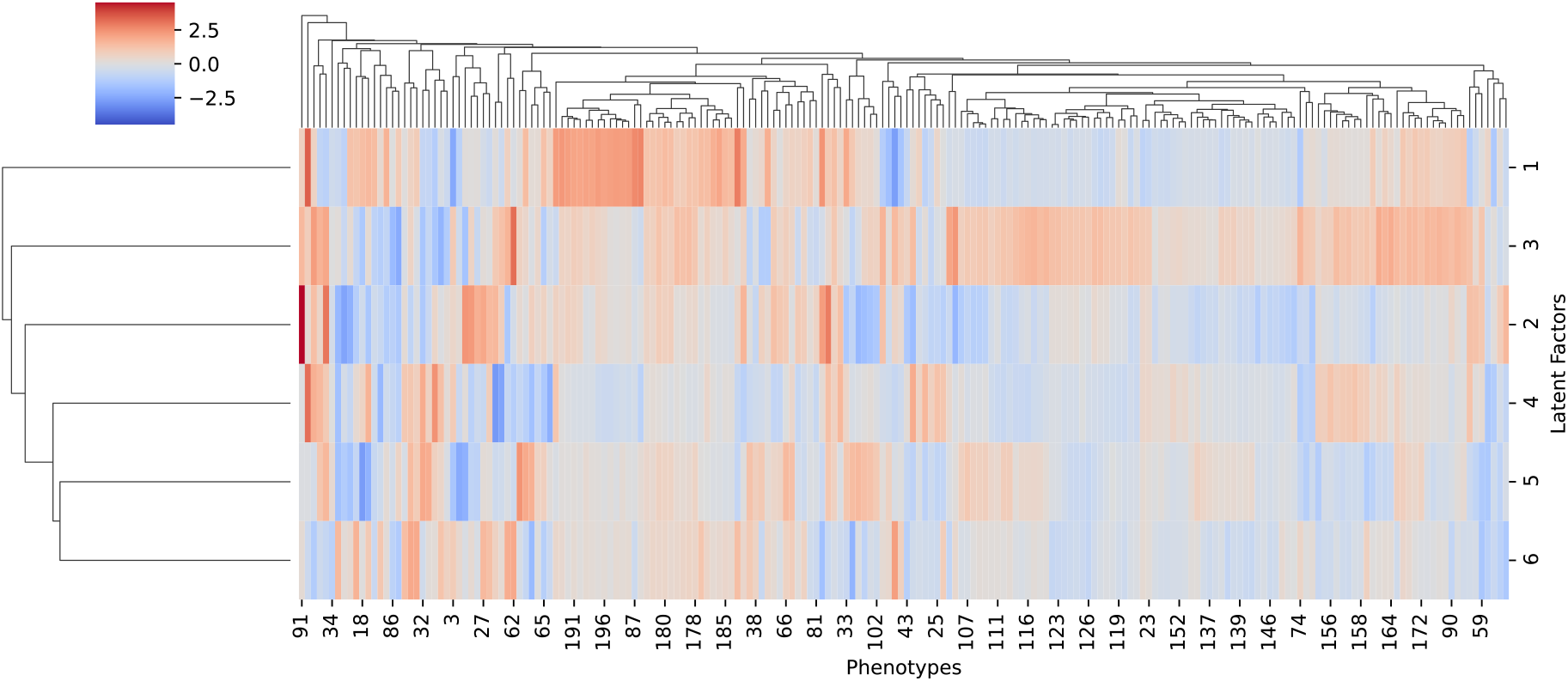
Clustermap of FABOr Normal Question Loadings on Simulated Binary and Ordinal Phenotype Datasets. Phenotypes are clustered based on the true **V** from the datasets, while latent factors are clustered based on FABOr’s estimation of **V**. Phenotypes 1 – 100 are binary, while phenotypes 101 – 200 are ordinal. Latent factors are numbered in descending order by the standard deviation of the question loadings and corrected for scale invariance by multiplying each latent factor by the standard deviation of the corresponding individual loadings.

When comparing the different variants of FABOr (Normal, Dirichlet-Beta, Dirichlet-Normal, and Lognormal), the Normal model performed slightly better across all simulated phenotypes, on both imputation accuracy to predict the observed values (by up to 5%) and MSE on **W** to recover the true latent factors (Table S1). While part of this could be due to the Gaussian distribution being the best prior to describe the data, it could also be due to the relative simplicity of optimizing Gaussian latent variables with variational inference compared to more complex probability distributions, or due to the underlying data generation scheme for the matrices following a true Gaussian distribution. For binary phenotypes, the Dirichlet-Beta and Dirichlet-Normal models perform slightly better than the Lognormal model, while for ordinal phenotypes the Lognormal model performs slightly better. In addition, whereas the Normal model converges at *K* = 7 for the binary dataset and *K* = 6 for the ordinal dataset, the non-Gaussian variants continue to achieve incremental gains in performance as the value of *K* approaches 10 (Fig. S2). Compared to the benchmarks, the non-Gaussian model variants perform better than *k*-Greedy BMF on the binary dataset and Ordinal MF/OrdNMF on the ordinal dataset, but they do not exceed the performance of Logistic MF on the binary dataset.

The accuracy of the base FABOr models and the benchmark models does not change significantly between the Full, MAR, and MNAR data missingness configurations, except for OrdNMF where the accuracy drops by ∼ 40% in the MAR and MNAR configurations. We believe the low performance of OrdNMF is due to two design choices. First, OrdNMF is the only benchmark method in which the ordinal cutpoints are defined uniformly across the entire dataset instead of separately for each phenotype, which leads to low performance on all data missingness configurations. Second, OrdNMF is also the only benchmark method in which observed values are encoded starting from 1 instead of 0; to account for this, the cutpoints include an extra value to model the missingness threshold, but it is again being modeled for the entire dataset when it actually varies between each phenotype, subsequently leading to additional performance drops in the MAR and MNAR configurations.

The MNAR by Data extension creates only marginal gains or losses in accuracy for all FABOr models tested, and the MNAR by Model extension for the Normal model marginally worsens performance across all values of *K* tested (Fig. S3–S6). This indicates that for the current levels of missingness tested (∼ 20% on average, up to 35% for a single phenotype in the MNAR data configuration) the observed data points are sufficient to achieve a reasonable fit to the data. When comparing the average runtime of FABOr against the benchmark models (Table S2), all variants run comparable to Logistic MF, slightly faster than Ordinal MF and OrdNMF, and significantly faster than *k*-Greedy BMF, even while incorporating Bayesian priors and additional missingness information which the benchmark methods do not include.

### 4.2 Application to Real-World Questionnaire Data

On the real-world questionnaires from the SPARK dataset, the base Normal model from FABOr performs 2% better compared to Logistic MF and 5% better than *k*-Greedy BMF on the SCQ for the best-performing value of *K*, and it performs 4% better than Ordinal MF and 23% better than OrdNMF on RBS-R. In particular, while the accuracy of the benchmark models decreases as *K* increases, the accuracy of FABOr increases, indicating that the model is able to adjust and use additional latent factors to capture the relationships in the data (Fig. S7). Most of the other observations noted on the simulated phenotypes hold true for the real-world phenotypes, except that all non-Gaussian model variants of FABOr exceed Logistic MF for binary phenotypes on the SCQ, and the difference in accuracy is within 2% of the Normal model (Fig. S8), which suggests that the non-Gaussian variants are equally suited to model the more complex posterior distributions that are observed in real-world data compared to the Normal model.

In the question loadings for the best value of *K* with FABOr, all variants show an underlying correlation between the first 20 questions and last 20 questions in the SCQ (Fig. 3). This correlation appears to map to the first 20 questions being related to verbal abilities and the last 20 questions centering on non-verbal abilities and interactions in social contexts with multiple people, despite no pre-defined structure or grouping in the questionnaire beyond the order of the questions. For RBS-R (Fig. 4), all variants show a weak to moderate correlation between the first 15 questions, which corresponds with the Stereotyped Behavior and Self-Injurious Behavior subscales of the questionnaire. In both datasets, most of the signal strength is concentrated in the first two latent factors, though both Dirichlet-Beta and Dirichlet-Normal show higher signal in the later latent factors.

**Fig. 3:**
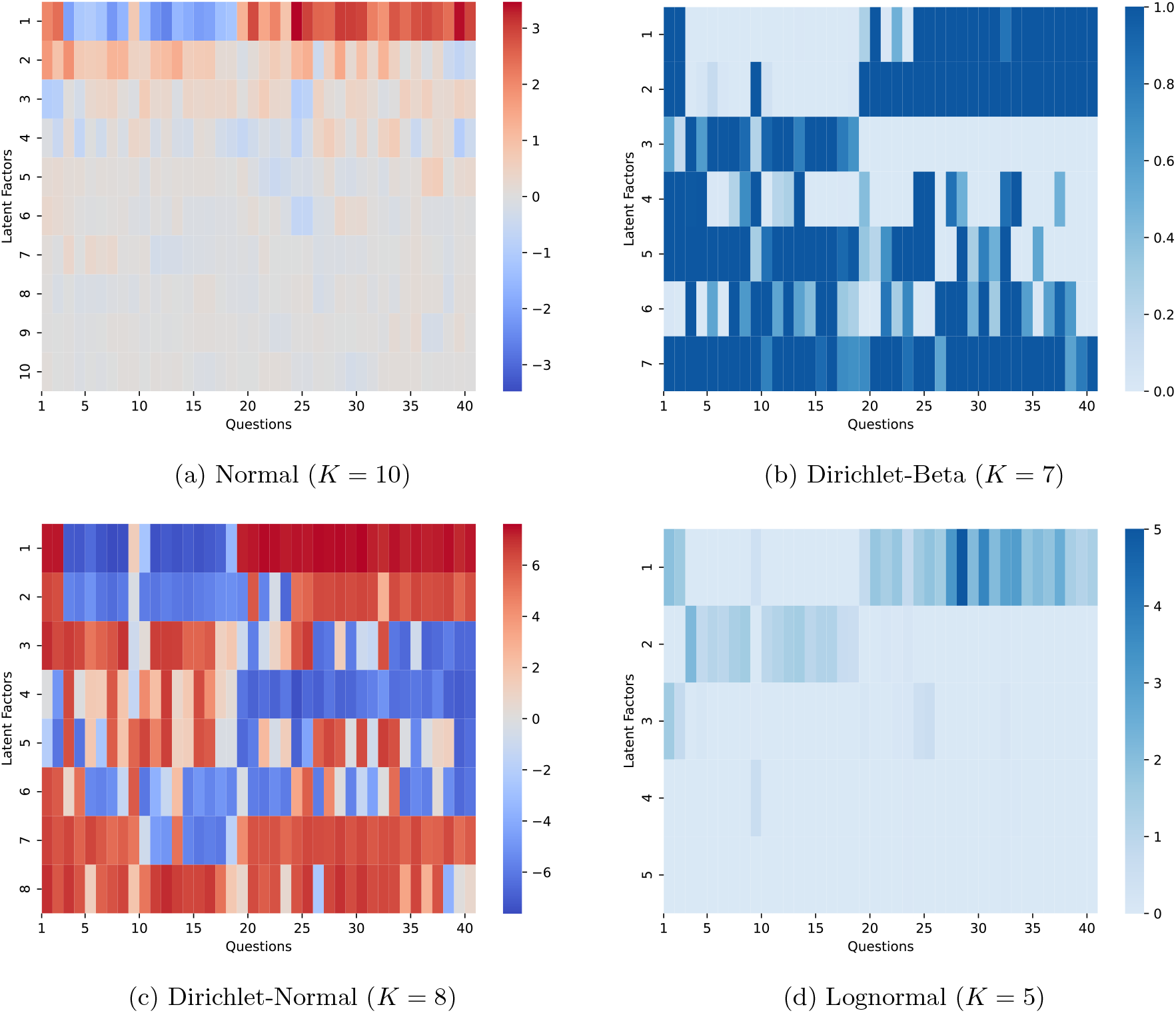
Heatmap of FABOr Question Loadings on the Social Communication Questionnaire (SCQ) from the Simons Foundation SPARK dataset. All latent factors are sorted in descending order by the standard deviation of the question loadings. Normal and Lognormal are corrected for scale invariance by multiplying each latent factor by the standard deviation of the corresponding individual loadings.

**Fig. 4:**
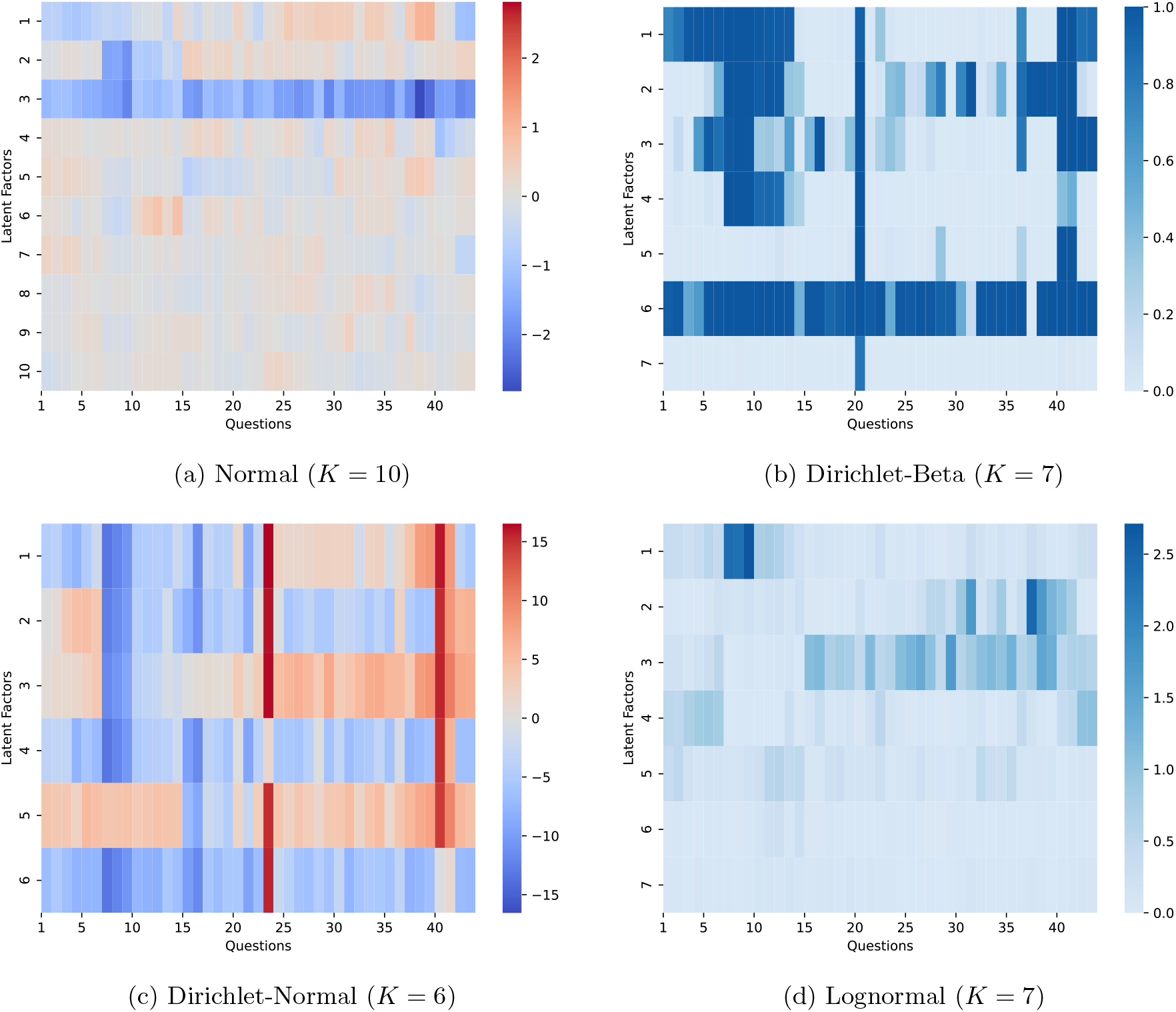
Heatmap of FABOr Question Loadings on the Repetitive Behavior Scale-Revised (RBS-R) from the SPARK dataset. All latent factors are sorted in descending order by the standard deviation of the question loadings. Normal and Lognormal are corrected for scale invariance by multiplying each latent factor by the standard deviation of the corresponding individual loadings.

In the individual loadings from FABOr (Fig. 5), all variants show underlying differences between ASD cases and Non-ASD controls. In the Normal and Lognormal models, ASD cases and Non-ASD controls form overlapping clusters in the latent space of the top 2 latent factors for the SCQ. The overall variance of ASD cases is ∼ 22% larger for the Normal model and ∼ 50% larger for the Lognormal model compared to Non-ASD controls. For the RBS-R which was only completed by ASD cases, the spread of the Lognormal model in the latent space is slightly more asymmetric compared to the Normal model. In the Dirichlet-Beta and Dirichlet-Normal models, the Non-ASD controls are clustered towards one corner of the simplex while the ASD cases fill the rest of the latent space. In both the SCQ and the RBS-R, individuals are highly concentrated towards the corners and edges of the simplex, suggesting that the models are identifying sparse latent factors.

**Fig. 5:**
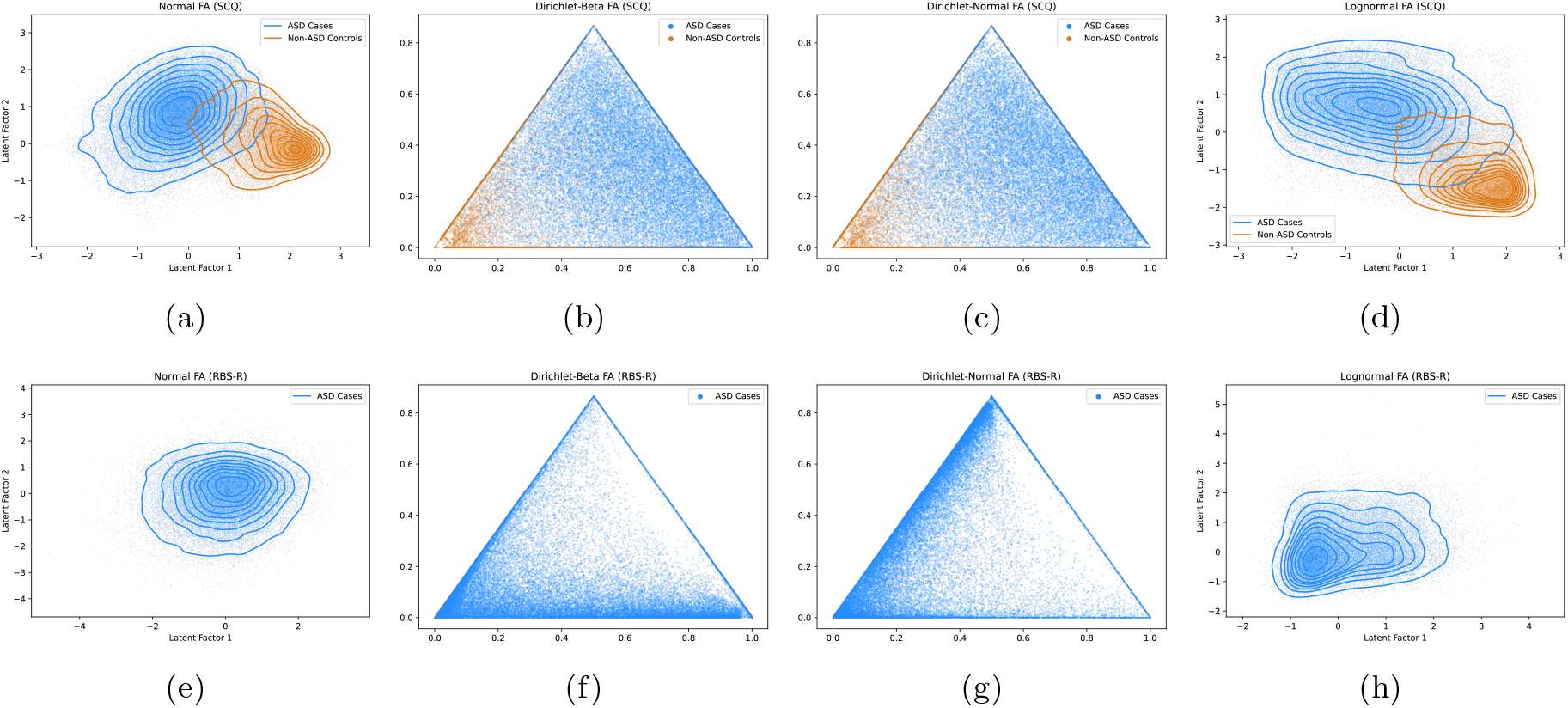
Latent Space of FABOr Individual Loadings on the SPARK dataset. (a)–(d) Latent Factors in SCQ for Normal (a), Dirichlet-Beta (b), Dirichlet-Normal (c), and Lognormal (d) models. (e)–(h) Latent Factors in RBS-R for Normal (e), Dirichlet-Beta (f), Dirichlet-Normal (g), and Lognormal (h) models. Normal and Lognormal loadings are shown with the top 2 latent factors at the best-performing value of *K*. Dirichlet-Beta and Dirichlet-Normal are shown on the simplex for *K* = 3. Lognormal loadings are log-adjusted.

## 5 Discussion

In this paper, we presented FABOr, a Bayesian framework for matrix factorization of binary and ordinal data that can be used to analyze the underlying relationships between clinical and self-reported phenotypes that fit within these constraints. It achieves this by using a Bernoulli likelihood to model binary phenotypes and a cutpoint-informed categorical likelihood to model ordinal phenotypes, while maintaining the flexibility for modeling the latent factors and low-rank matrices using various continuous prior distributions. Compared to current factor analysis methods available for binary and ordinal data, we showed that FABOr performed similar to better on simulated phenotypes and uniformly better on real-world phenotypes from the SPARK dataset. We also tested different prior distributions for the low-rank matrices in FABOr and found that Gaussian priors performed best, though there may be certain cases where a non-Gaussian prior is more desirable for interpretability of the latent factors. While we designed extensions of FABOr to handle MNAR data, we found that this did not lead to significant improvements in performance for most of the phenotypes we tested, but we anticipate they may be more useful for sparse datasets that have higher rates of missingness (e.g., above 50%).

The latent factors from FABOr can be treated as additional phenotypes via the individual loadings to supplement the original data, which can be useful especially when working with datasets that have self-reported phenotypes with varying degrees of accuracy [18]. The distribution of values for each latent factor is influenced by the selection of prior for **U** (Fig. 6): Normal and Lognormal latent factors can be interpreted as quantitative phenotypes, while Dirichlet-Beta and Dirichlet-Normal latent factors can be interpreted as binary phenotypes. These latent factors could be tested for genotypic associations like any other phenotypes via genome-wide association studies using single-variant or gene-based tests, and we are working on one such genome-wide association study using whole-exome sequencing and genotyping array data from the SPARK dataset.

**Fig. 6:**
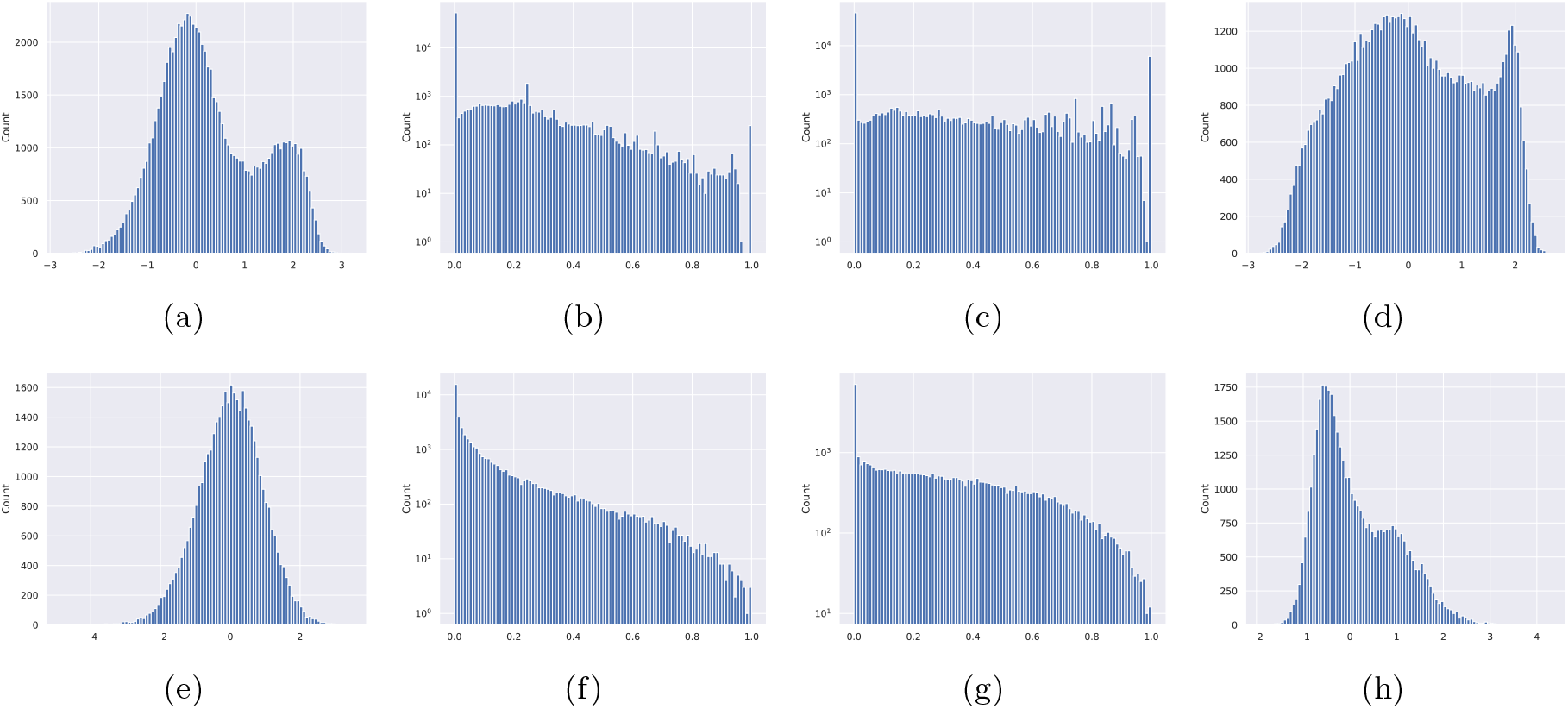
Distribution of FABOr Individual Loadings on the SPARK dataset. (a)–(d) Distribution of Latent Factor 1 in SCQ across individuals for Normal (a), Dirichlet-Beta (b), Dirichlet-Normal (c), and Lognormal (d) models. (e)–(h) Distribution of Latent Factor 1 in RBS-R across individuals for Normal (e), Dirichlet-Beta (f), Dirichlet-Normal (g), and Lognormal (h) models. Dirichlet-Beta and Dirichlet-Normal loadings are plotted on a log scale. Lognormal loadings are log-adjusted.

## 6 Acknowledgments

This research was supported in part by NSF grants DGE-2036197 (N.S.) and DBI-2146398 (D.A.K.), NIH grants F31DC022183 (N.S.) and R01MH130879 (D.A.K.), and the Columbia University SIRS Blavatnik Acceleration Fund (D.A.K.).

## 7 Author Contributions

N.S. and D.A.K. designed the research. N.S. designed the Bayesian factor analysis models and phenotype generation algorithms with input from D.A.K. N.S. implemented the models and performed all experiments and analyses. N.S. and D.A.K. wrote the manuscript.

---

**Table S1:**
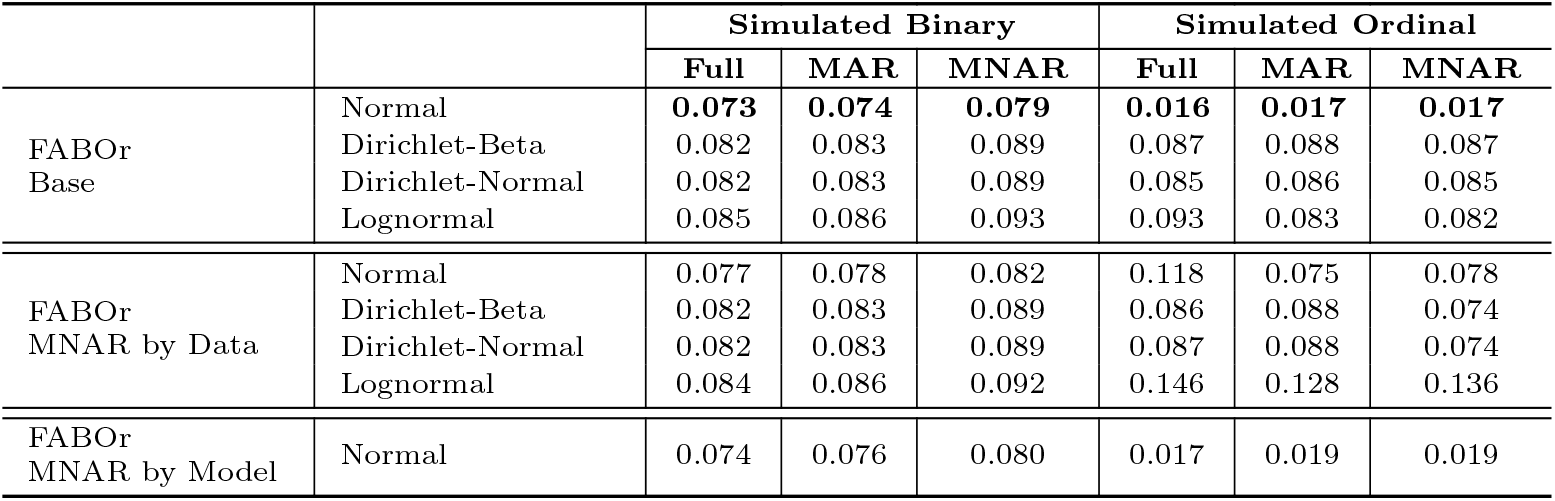
Evaluation of latent factor recovery via **W** on simulated phenotypes. The average mean squared error (MSE) across 10 random restarts is reported for the best performing value of *K*.

**Table S2:** Average runtime of models (in seconds) across 10 random restarts on simulated and real-world phenotypes for the best performing value of *K*. Experiments were performed on an NVIDIA RTX Pro 6000 Blackwell GPU.

|  |  | Simulated Binary |  |  | Simulated Ordinal |  |  | Real-World |  |
| --- | --- | --- | --- | --- | --- | --- | --- | --- | --- |
|  |  | Full | MAR | MNAR | Full | MAR | MNAR | SCQ | RBS-R |
| Benchmarks | Logistic MF | 5.844 | **3.681** | **3.661** | — | — | — | **1.357** | — |
| | $k$-Greedy BMF [9] | 775.1 | 779.3 | 789.8 | — | — | — | 630.1 | — |
|  | Ordinal MF | — | — | — | 36.28 | 31.29 | 31.61 | — | **3.210** |
|  | OrdNMF [6] | — | — | — | 24.98 | 18.62 | 32.65 | — | 6.942 |
| FABOr<br>Base | Normal | 6.670 | 5.415 | 5.431 | 16.44 | 16.50 | 16.30 | 4.899 | 5.310 |
|  | Dirichlet-Beta | 5.562 | 5.543 | 5.554 | 16.31 | **16.20** | 16.79 | 4.146 | 5.620 |
|  | Dirichlet-Normal | **5.342** | 5.349 | 5.337 | **16.30** | 16.40 | **16.01** | 3.787 | 5.349 |
|  | Lognormal | 5.899 | 5.883 | 5.869 | 17.29 | 17.24 | 17.19 | 4.193 | 5.737 |
| FABOr<br>MNAR by Data | Normal | 9.264 | 9.284 | 9.270 | 19.45 | 19.70 | 19.31 | 4.473 | 6.280 |
|  | Dirichlet-Beta | 10.28 | 10.26 | 10.26 | 19.69 | 19.84 | 19.86 | 4.652 | 6.594 |
|  | Dirichlet-Normal | 9.158 | 9.244 | 9.216 | 19.04 | 19.65 | 19.62 | 4.415 | 6.339 |
|  | Lognormal | 10.57 | 10.62 | 10.61 | 20.44 | 20.58 | 20.74 | 4.955 | 6.717 |
| FABOr<br>MNAR by Model | Normal | 9.882 | 9.854 | 9.888 | 19.69 | 19.91 | 20.02 | 6.888 | 8.468 |

**Fig. S1:**
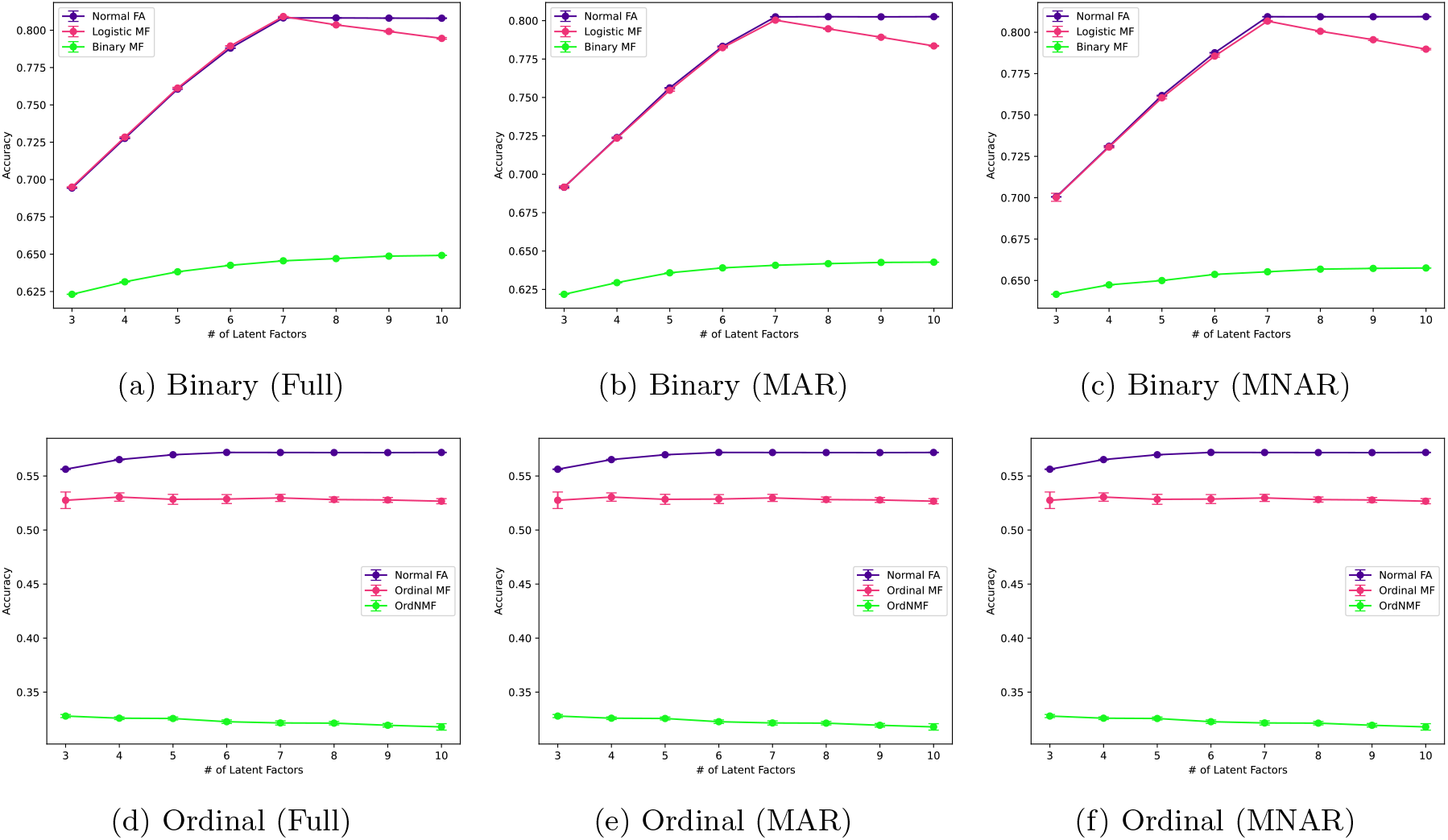
Comparison of FABOr and Model Benchmarks on Simulated Phenotypes. Purple lines are FABOr models, magenta lines are classical matrix factorization models with binary/ordinal likelihoods, and green lines are specialized binary/ordinal matrix factorization models. Mean ± standard deviation of model accuracy reported across 10 restarts (i.e., random initializations).

**Fig. S2:**
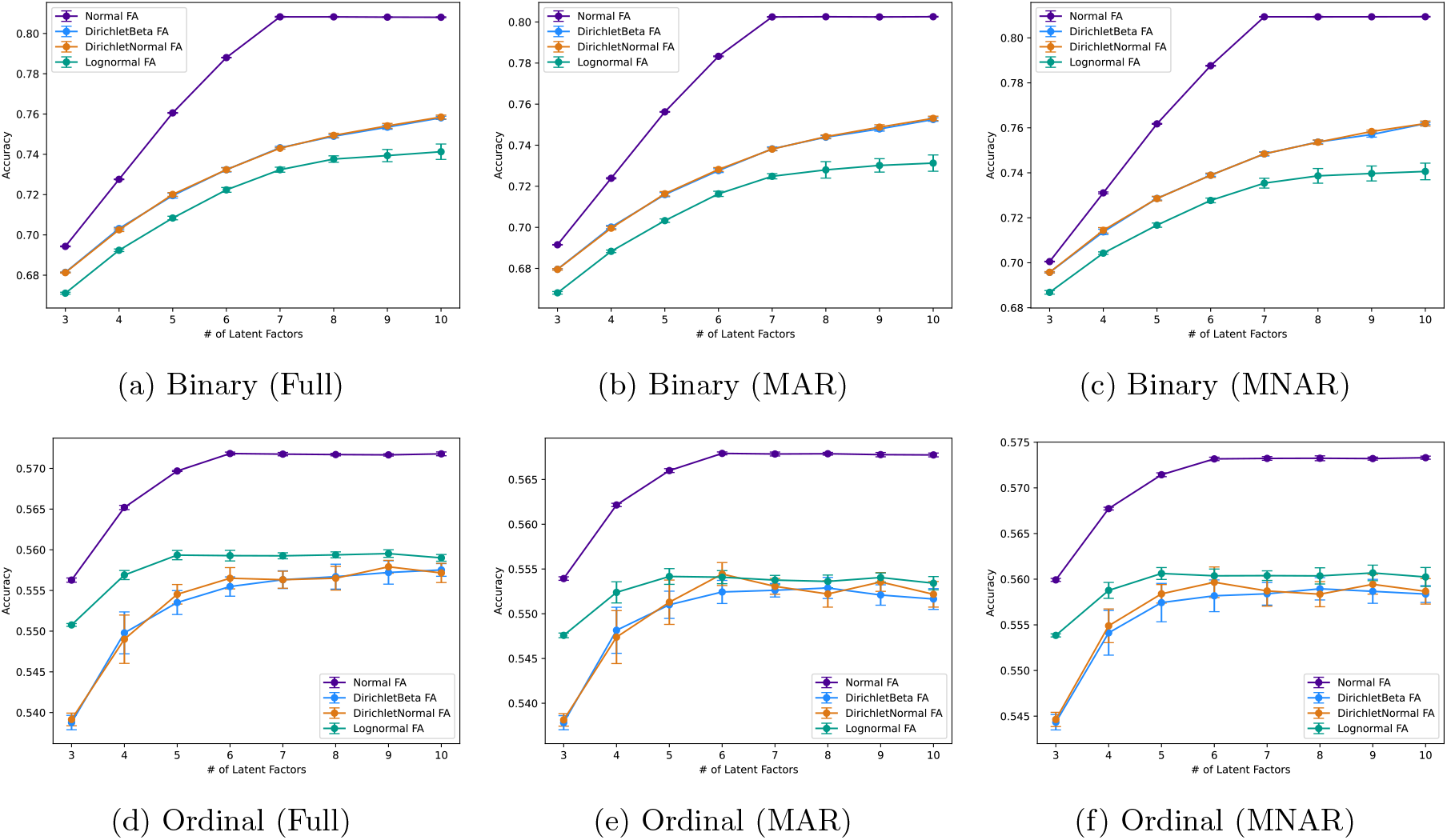
Comparison of FABOr Variants on Simulated Phenotypes. Mean ± standard deviation of model accuracy reported across 10 restarts (i.e., random initializations).

**Fig. S3:**
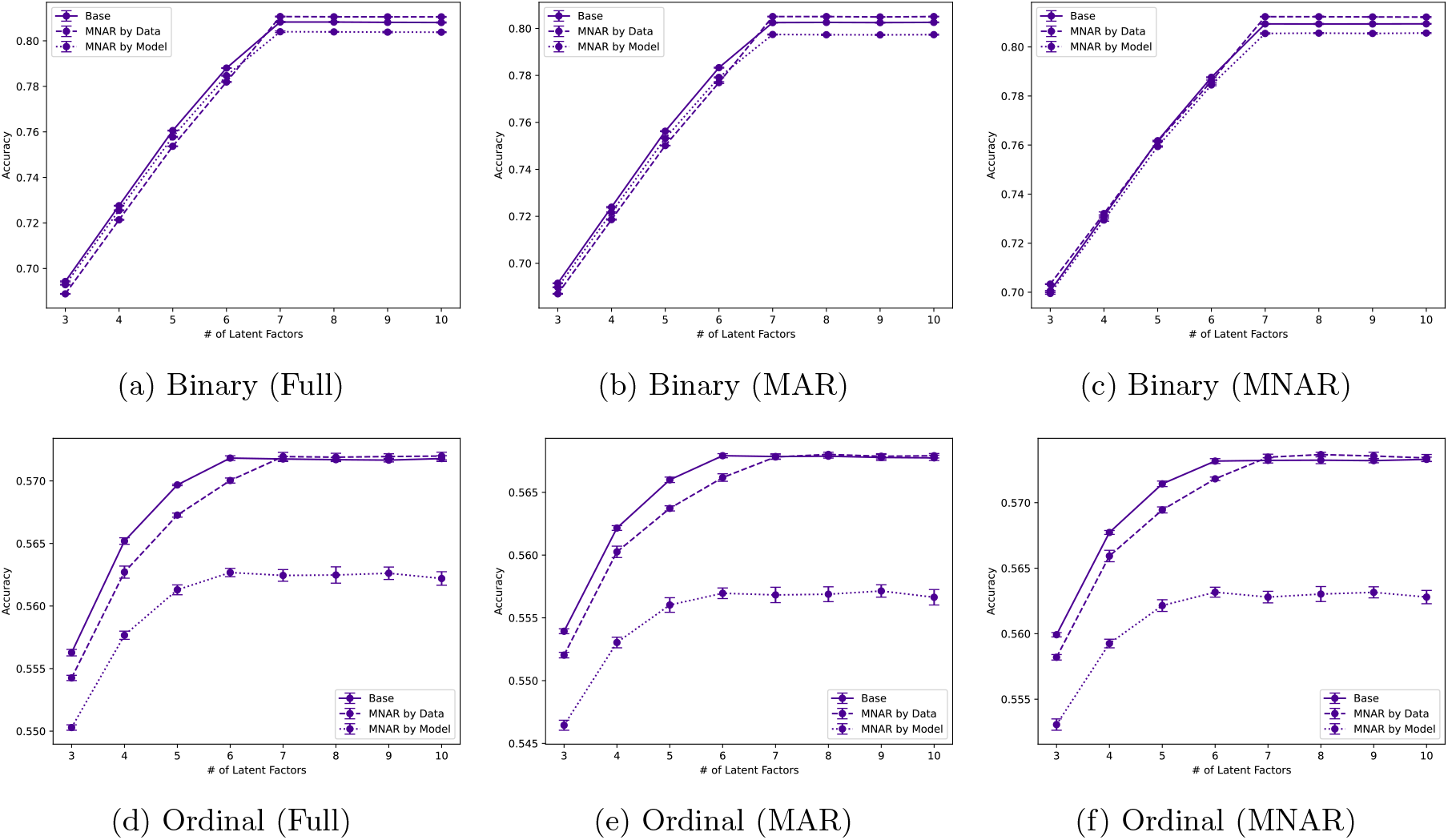
Comparison of FABOr Normal with MNAR Extensions on Simulated Phenotypes. Mean ± standard deviation of model accuracy reported across 10 restarts (i.e., random initializations).

**Fig. S4:**
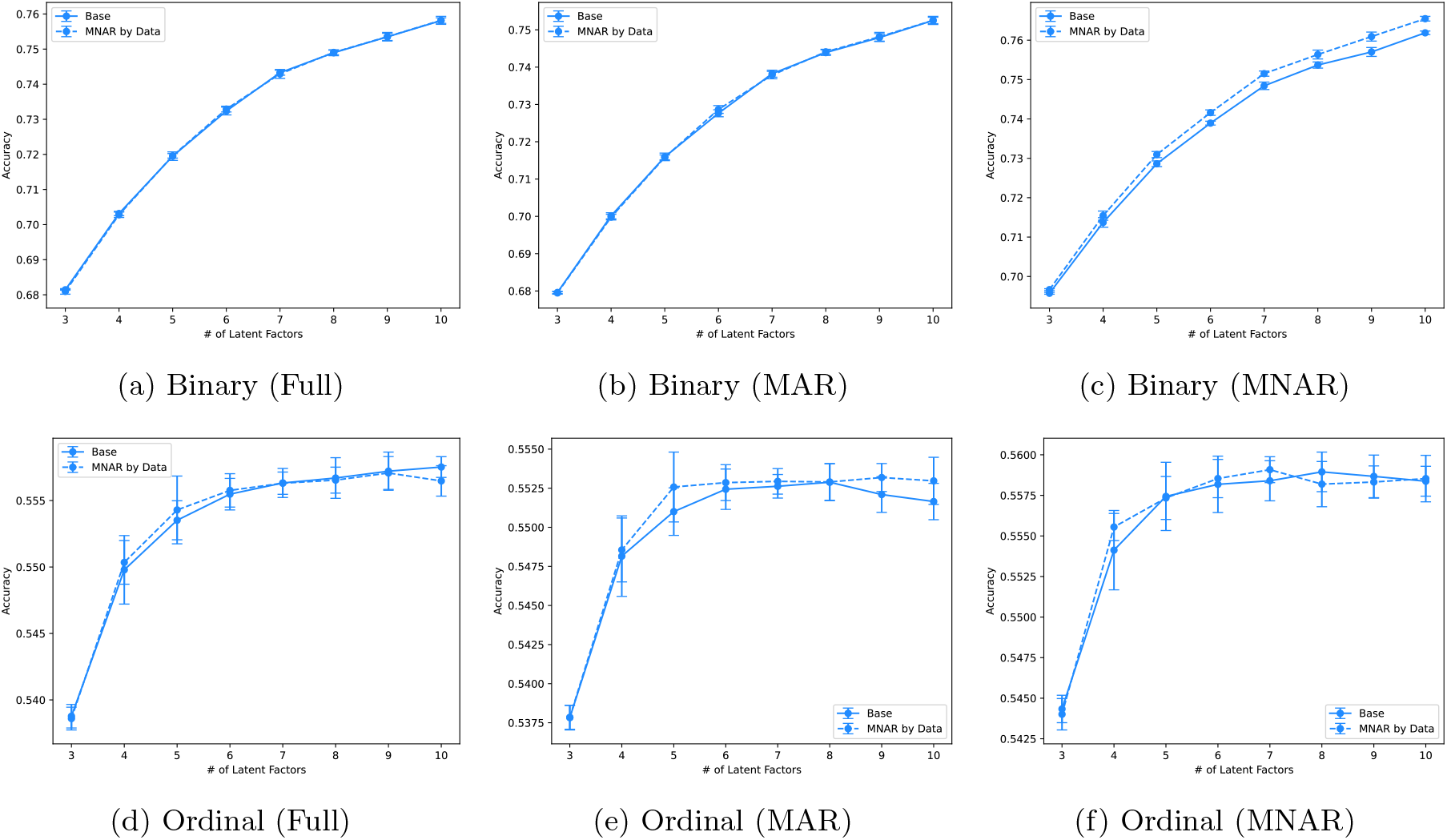
Comparison of FABOr Dirichlet-Beta with MNAR Extensions on Simulated Phenotypes. Mean ± standard deviation of model accuracy reported across 10 restarts (i.e., random initializations).

**Fig. S5:**
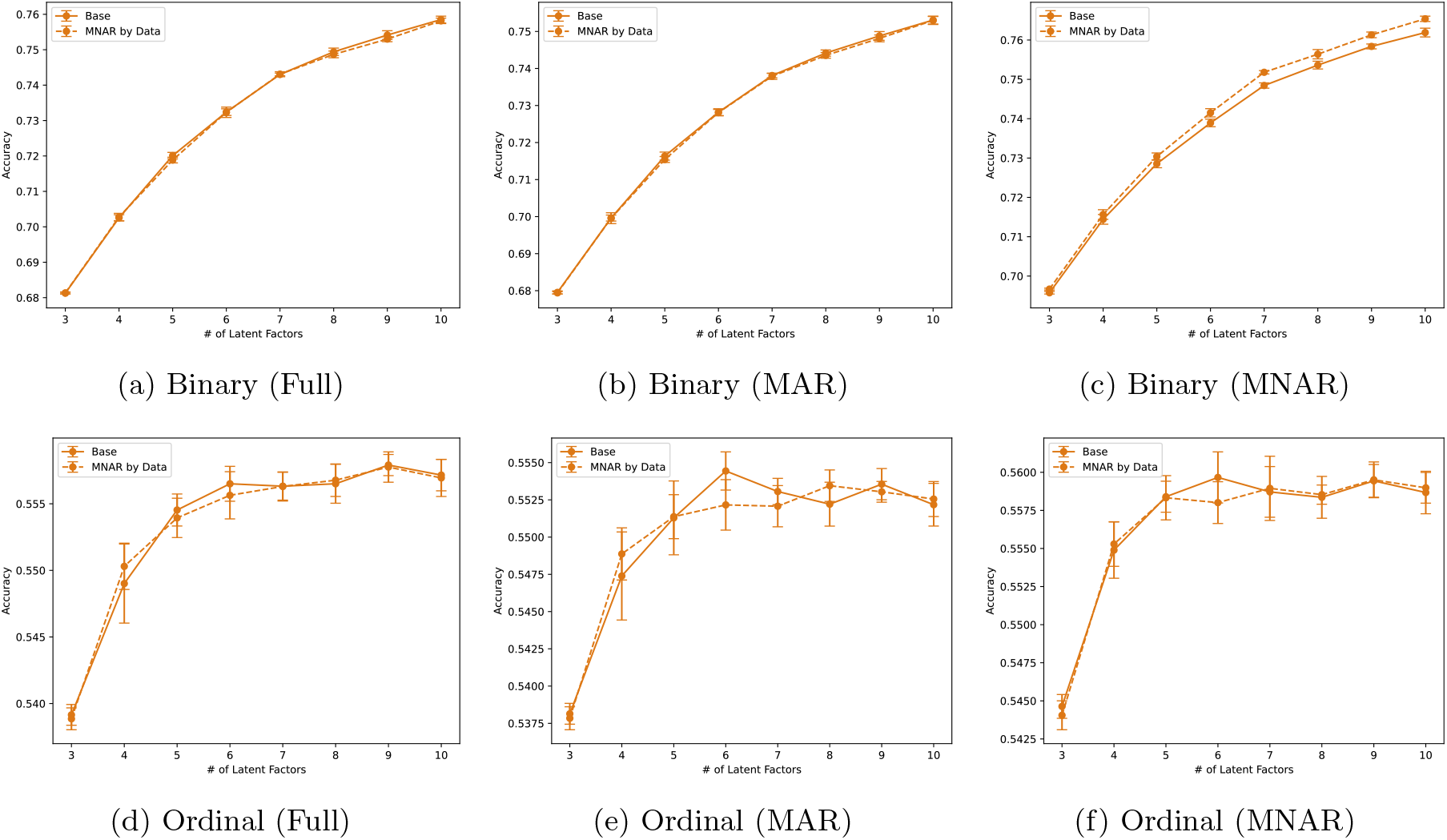
Comparison of FABOr Dirichlet-Normal with MNAR Extensions on Simulated Phenotypes. Mean ± standard deviation of model accuracy reported across 10 restarts (i.e., random initializations).

**Fig. S6:**
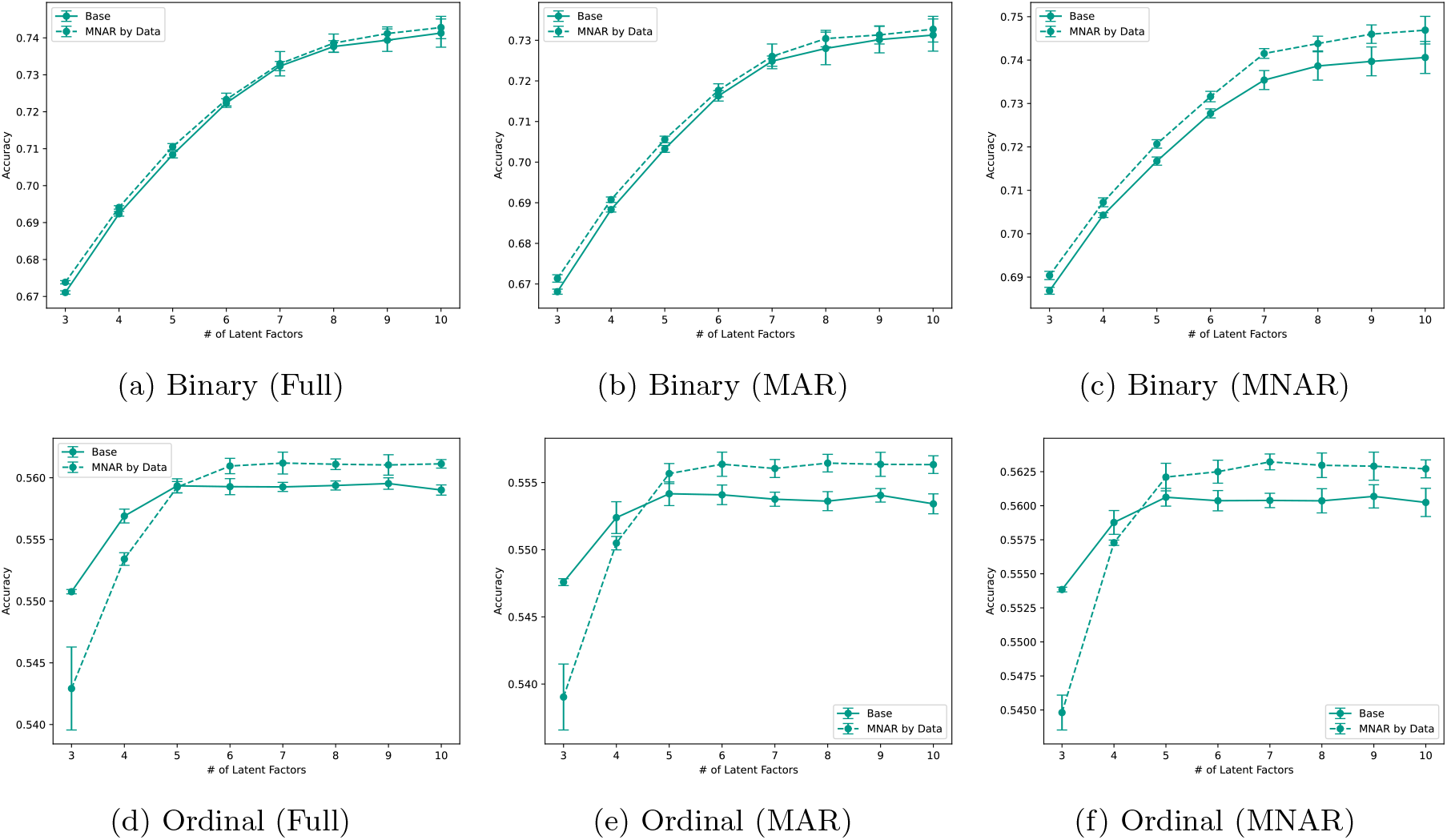
Comparison of FABOr Lognormal with MNAR Extensions on Simulated Phenotypes. Mean ± standard deviation of model accuracy reported across 10 restarts (i.e., random initializations).

**Fig. S7:**
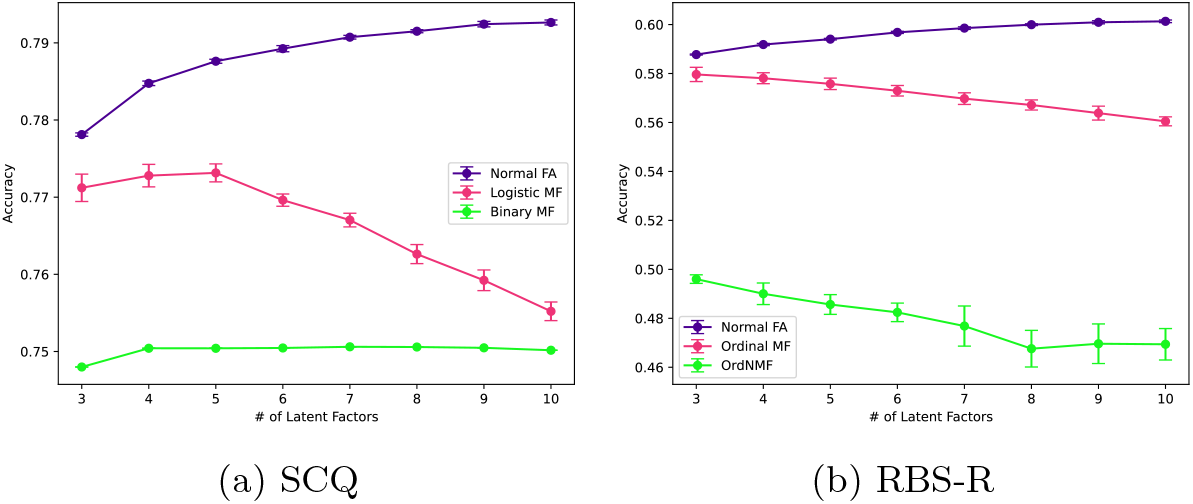
Comparison of FABOr and Model Benchmarks on Real-World Phenotypes from the SPARK dataset. Purple lines are FABOr models, magenta lines are classical matrix factorization models with binary/ordinal likelihoods, and green lines are specialized binary/ordinal matrix factorization models. Mean ± standard deviation of model accuracy reported across 10 restarts (i.e., random initializations).

**Fig. S8:**
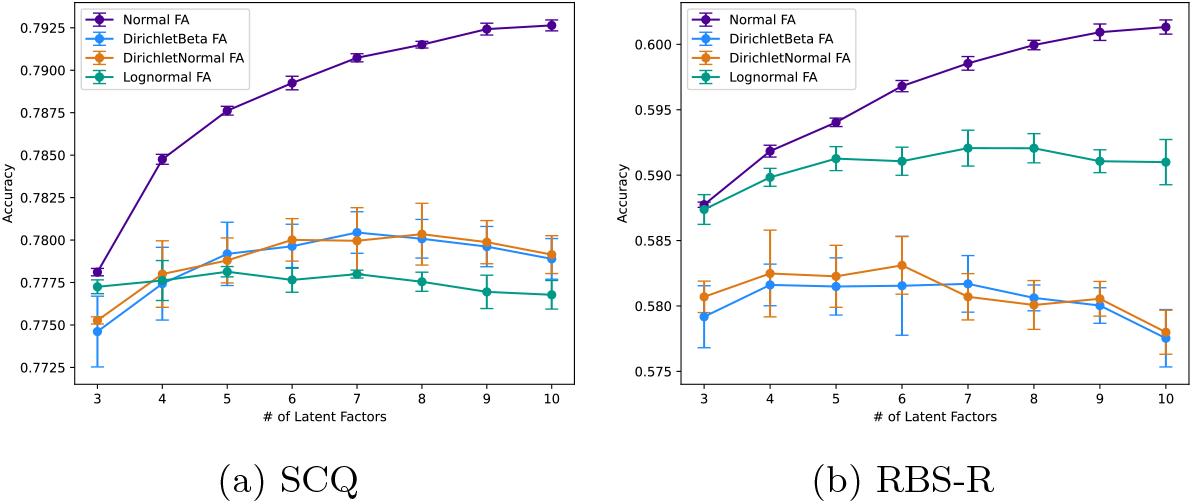
Comparison of FABOr Variants on Real-World Phenotypes from the SPARK dataset. Mean ± standard deviation of model accuracy reported across 10 restarts (i.e., random initializations).

## Notes

### Competing Interest Statement

The authors have declared no competing interest.

